# Natural variation in the sequestosome-related gene, *sqst-5*, underlies zinc homeostasis in *Caenorhabditis elegans*

**DOI:** 10.1101/2020.07.10.196857

**Authors:** Kathryn S. Evans, Stefan Zdraljevic, Lewis Stevens, Kimberly Collins, Robyn E. Tanny, Erik C. Andersen

## Abstract

Zinc is an essential trace element that acts as a co-factor for many enzymes and transcription factors required for cellular growth and development. Altering intracellular zinc levels can produce dramatic effects ranging from cell proliferation to cell death. To avoid such fates, cells have evolved mechanisms to handle both an excess and a deficiency of zinc. Zinc homeostasis is largely maintained via zinc transporters, permeable channels, and other zinc-binding proteins. Variation in these proteins might affect their ability to interact with zinc, leading to either increased sensitivity or resistance to natural zinc fluctuations in the environment. We can leverage the power of the roundworm nematode *Caenorhabditis elegans* as a tractable metazoan model for quantitative genetics to identify genes that could underlie variation in responses to zinc. We found that the laboratory-adapted strain (N2) is resistant and a natural isolate from Hawaii (CB4856) is sensitive to micromolar amounts of exogenous zinc supplementation. Using a panel of recombinant inbred lines, we identified two large-effect quantitative trait loci (QTL) on the left arm of chromosome III and the center of chromosome V that are associated with zinc responses. We validated and refined both QTL using near-isogenic lines (NILs) and identified a naturally occurring deletion in *sqst-5*, a sequestosome-related gene, that is associated with resistance to high exogenous zinc. We found that this deletion is relatively common across strains within the species and that variation in *sqst-5* is associated with zinc resistance. Our results offer a possible mechanism for how organisms can respond to naturally high levels of zinc in the environment and how zinc homeostasis varies among individuals.

**Author summary:** Zinc, although an essential metal, can be toxic if organisms are exposed to concentrations that are too high or too low. To prevent toxicity, organisms have evolved mechanisms to regulate zinc uptake from the environment. Here, we leveraged genetic variation between two strains of the roundworm *Caenorhabditis elegans* with different responses to high exogenous zinc to identify genes that might be involved in maintaining proper zinc levels. We identified four loci that contributed to differential zinc responses. One of these loci was the sequestosome-related gene *sqst-5*. We discovered that targeted deletions of *sqst-5* caused an increase in resistance to zinc. Although SQST-5 contains a conserved zinc-binding protein domain, it has yet to be directly implicated in the *C. elegans* zinc response pathway. We identified two common forms of genetic variation in *sqst-5* among 328 distinct strains, suggesting that variation in *sqst-5* must have emerged multiple times, perhaps in response to an environment of high zinc. Overall, our study suggests a natural context for the evolution of zinc response mechanisms.

## Introduction

Heavy metals such as zinc, iron, and copper are known to play important roles in many biological systems [1,2]. Of these metals, zinc is the most abundant and is essential for proper function of many proteins, including enzymes and transcription factors [3]. In addition to its function as a cofactor, zinc can act as a signaling molecule in neurons [4–7] and is known to play a role in cell-fate determination [8–12]. Because of its many functions, zinc deficiency has been shown to cause major defects, including growth inhibition and death in several species [8,13–16]. On the other hand, excess zinc can also be toxic, displaying phenotypic effects similar to copper deficiency, anemia, and neutropenia [17]. Although the exact mechanisms are unknown, the data suggest that excess zinc might bind ectopically to other proteins, displacing similar metals such as copper or magnesium from these proteins [8]. Because of this need for intracellular zinc balance even though environmental zinc might fluctuate, biological systems must use proper zinc homeostasis mechanisms for uptake, distribution, efflux, and detoxification [13].

The nematode *Caenorhabditis elegans* is a tractable metazoan model for studying the molecular mechanisms of zinc homeostasis and toxicity [8,18–20]. As observed in other organisms, zinc is essential for *C. elegans* growth [21]. In fact, it is estimated that about 8% of the *C. elegans* genes (1,600 genes) encode zinc-binding proteins [22]. However, zinc is also toxic to the nematode at higher concentrations [21]. High exogenous zinc can cause several defects including decreased growth rate and survival, suppression of the multivulva phenotype, and formation of bilobed lysosome-related organelles in intestinal cells [8]. Genetic screens have identified several genes that act to increase sensitivity to high levels of zinc (*haly-1, natc-1*, and *daf-21*) [8,23–26]. However, mutations in these genes cause a change in response to multiple stressors (including metals, heat, and oxidation), suggesting they are not specific to zinc homeostasis [8,25]. In addition to these nonspecific zinc proteins, *C. elegans* also has two complementary families (composed of 14 proteins each) of zinc transporters responsible for maintaining constant intracellular zinc concentrations via import and export [8]. Four of these zinc exporters (*cdf-1, cdf-2, sur-7*, and *ttm-1*) have been shown to promote resistance to high zinc toxicity [8,9,21,23,27,28].

Although much is already known about zinc biology in *C. elegans*, previous studies were performed using a single laboratory-adapted strain (N2) that is known to differ significantly, both genetically and phenotypically, from wild isolates in the species [29]. As a complementary approach, we can leverage the power of natural genetic diversity among wild isolates [30–32] to identify novel mechanisms of zinc homeostasis and gain insights into the evolution of this process. We used a large panel of recombinant inbred advanced intercross lines (RIAILs) [33–35] constructed from a multi-generational cross between two genetically and phenotypically diverged strains, N2, the laboratory-adapted strain, and CB4856, a wild isolate from Hawaii [36]. This panel of RIAILs has been leveraged in several linkage mapping analyses, identifying hundreds of quantitative trait loci (QTL) [35,37–63]. In combination with a high-throughput phenotyping assay to measure animal length, optical density, and brood size [34], several quantitative trait genes (QTG) [35,63] and quantitative trait nucleotides (QTN) [41,42] underlying fitness-related traits have been described.

Here, we use linkage mapping analysis to identify four QTL in response to high exogenous zinc. Several genes previously identified to be involved in the zinc response were found within the QTL on chromosomes V and X. However, no known zinc-related genes were located in the large-effect chromosome III QTL, suggesting a potentially novel mechanism of zinc homeostasis. We constructed reciprocal near-isogenic lines (NILs) for each QTL and used them to validate the two large-effect QTL on chromosomes III and V. Expression QTL mapping and mediation analysis identified a single candidate gene, *sqst-5*, with predicted zinc ion-binding capability. We used CRISPR-Cas9 genome editing to show that strains without *sqst-5* were significantly more resistant to zinc supplementation than strains with a functional copy of *sqst-5*, suggesting a new role for this gene in zinc regulation. In addition to CB4856, several other wild isolates were found to share a 111 bp deletion in *sqst-5*. Moreover, we identified a second group of strains with a distinct haplotype of variation at *sqst-5* that was also associated with zinc resistance. Together, these data suggest that the regulation of zinc in nematodes is complex, but binding and accumulation of excess zinc might be a mechanism to respond to exogenous zinc.

## Results

### Natural genetic variation in response to zinc is complex

We exposed four genetically divergent strains of *C. elegans* (N2, CB4856, JU258, and DL238) to increasing concentrations of exogenous zinc and measured their development (animal length, optical density, and normalized optical density) and reproductive ability (brood size) with a high-throughput assay using the COPAS BIOSORT (see Methods) [34,35,41–43]. In the presence of high concentrations of zinc, animals of all strains had smaller broods, shorter lengths, and were less optically dense compared to non-treated animals (**S1 Fig and S1 File**). Because nematodes grow longer and become more optically dense as they develop, these results suggest a zinc-induced developmental delay. Furthermore, the lower brood size of animals treated with zinc suggest that exogenous zinc hinders reproductive ability in some way. In addition to these overall trends, we also observed significant phenotypic variation among strains. For example, although all strains had smaller lengths in the presence of exogenous zinc, animals of the N2 strain were the largest (most resistant to zinc), and animals of the CB4856 strain were smaller (more sensitive to zinc). At 500 μM zinc, a concentration that both maximizes among-strain and minimizes within-strain phenotypic variation, we identified a substantial heritable genetic component for two highly correlated developmental traits: animal length (*H*^2^ = 0.48, 95% CI [0.30, 0.61]) and optical density (*H*^2^ = 0.48, 95% CI [0.28, 0.59]) (**S2 File**).

To investigate the genetic basis of zinc response, we exposed a panel of 253 RIAILs derived from a cross between the N2 and CB4856 strains (set 2 RIAILs, see Methods) to high exogenous zinc (**S3 File**). In these conditions, the N2 animals were longer (**S2 Fig**) and more optically dense (**Fig 1A**) than the CB4856 animals, and were thus more resistant to high zinc supplementation. Interestingly, many of the RIAILs were either more resistant than N2 or more sensitive than CB4856, suggesting that loci of opposite genotypes are either acting additively or interacting in the RIAILs to produce the observed transgressive phenotypes [64]. Linkage mapping analysis identified 12 QTL across all traits, representing five unique QTL on chromosomes III, IV, V, and X (**S2 Fig and S4 File**). Because genetic architectures looked similar across these traits, we chose to focus our analyses on optical density to avoid redundant analyses of correlated traits (**Fig 1B**). Together, the four QTL underlying animal optical density explain 40.5% of the phenotypic variation among the RIAILs. As expected, QTL of opposite effects were observed. Strains with the CB4856 allele on chromosome III were more resistant to zinc than strains with the N2 allele at this locus. By contrast, strains with the CB4856 alleles on chromosomes IV, V, and X were more sensitive to zinc than strains with the N2 alleles at these loci (**Fig 1C, S3 and S4 Files**). We scanned the genome for interactions between pairs of genomic markers that might affect the phenotypic distribution of the RIAIL panel and identified no significant interactions (**S3 Fig and S5 File**). We further examined the additivity of the two QTL with the largest and opposite effect sizes (QTL on chromosomes III and V). We concluded that RIAILs with the CB4856 allele on chromosome III and the N2 allele on chromosome V were the most resistant, and RIAILs with the N2 allele on chromosome III and the CB4856 allele on chromosome V were the most sensitive (**S4 Fig and S3 File**). Furthermore, the effect size of the chromosome III locus was similar regardless of the genotype on chromosome V (**S4 Fig and S3 File**), and no significant interaction term was identified using a linear model (ANOVA, *p* = 0.251). These results suggest that multiple additive QTL rather than interacting loci affect animal optical densities in zinc.

**Fig 1.**
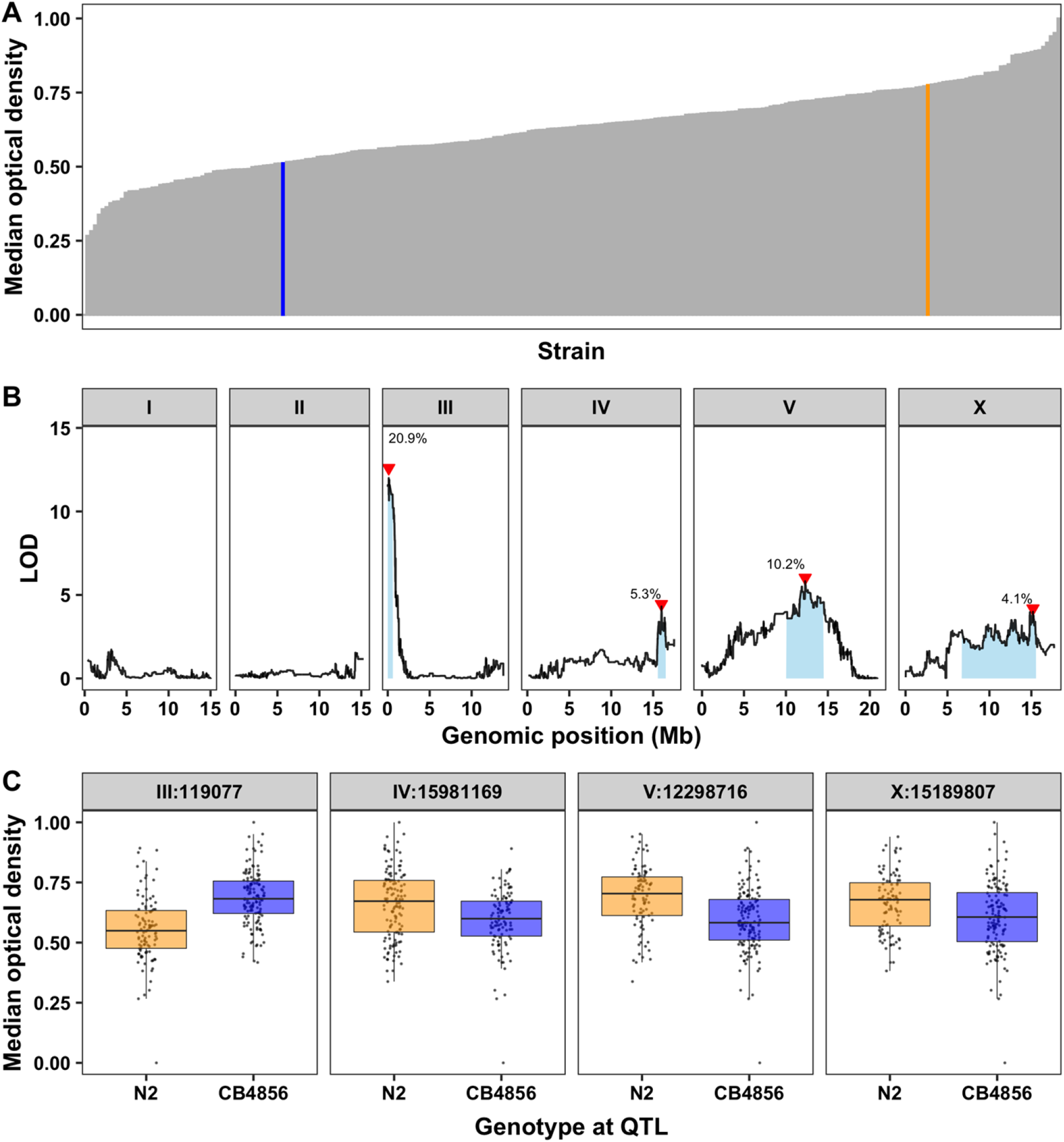
Linkage mapping identifies four QTL in response to high dietary zinc. **A)** Median optical densities (y-axis) of 253 RIAILs (x-axis) in response to zinc supplementation are shown. The parental strains are colored: N2, orange; CB4856, blue. **B)** Linkage mapping results for optical density (median.EXT) is shown. Genomic position (x-axis) is plotted against the logarithm of the odds (LOD) score (y-axis) for 13,003 genomic markers. Each significant QTL is indicated by a red triangle at the peak marker, and a blue rectangle shows the 95% confidence interval around the peak marker. The percentage of the total variance in the RIAIL population that can be explained by each QTL is shown above the QTL. **C)** For each QTL, the normalized residual median optical densities (y-axis) of RIAILs split by genotype at the marker with the maximum LOD score (x-axis) are plotted as Tukey box plots. Each point corresponds to a unique recombinant strain. Strains with the N2 allele are colored orange, and strains with the CB4856 allele are colored blue.

### Near-isogenic lines fractionate the chromosome V QTL into multiple additive loci

We first investigated whether any of the 28 known zinc transporters or any of the other 15 zinc-related genes [8] are located in one of the four detected QTL intervals. We discovered that three of these genes lie in the QTL on chromosome V and 11 lie in the QTL on chromosome X (**S6 File**). However, none of these zinc-related genes lie in either of the QTL on chromosomes III or IV (**S6 File**). To isolate and validate the effect of these four QTL, we constructed reciprocal near-isogenic lines (NILs) by introgressing a genomic region surrounding each of the QTL from the CB4856 strain into the N2 genetic background or vice versa (**S7 and S8 Files**). We then measured the animal optical densities in the presence of zinc for these strains to provide experimental evidence in support of each QTL independently. For the three QTL on chromosomes IV, V, and X, the N2 allele was associated with zinc resistance (**Fig 1C, S3 and S4 Files**). However, strains with the N2 allele crossed into a CB4856 genetic background on chromosomes IV and X were as sensitive as the CB4856 strain, and strains with the CB4856 allele crossed into an N2 genetic background were as resistant as the N2 strain (**S5 Fig, S9 and S10 Files**). These two QTL have the smallest effect sizes among the four QTL detected and each explain only 5% of the total phenotypic variation among the RIAILs. The lack of a significant difference between the NILs and their respective parental strains suggests that the QTL effect might be smaller than 5% and we were underpowered to detect the difference. Alternatively, the interval might contain QTL of opposing effects requiring additional smaller NILs.

By contrast, the NIL with the N2 allele surrounding the QTL on chromosome V introgressed into the CB4856 genetic background is significantly more resistant than the sensitive CB4856 strain (**S5 Fig, S9 and S10 Files**). This result confirms that genetic variation between the N2 and CB4856 strains on the center of chromosome V contributes to the differences in animal optical densities between the strains in the presence of high exogenous zinc. To further narrow this QTL, we created a panel of NILs with smaller regions of the N2 genome introgressed into the CB4856 genetic background. We exposed a subset of these NILs to zinc and measured their optical densities. We found strains with the resistant N2 phenotype (ECA481; V:9.6-13.8 Mb), strains with the sensitive CB4856 phenotype (ECA411; V:11.3-13.9 Mb), and strains with an intermediate phenotype (ECA437; V:10.5-13.8 Mb) (**Fig 2, S10 and S11 Files**). These data imply the existence of at least two loci (V:9.6-10.5 Mb and V:10.5-11.3 Mb) at which the N2 allele confers resistance to zinc. The intermediate strain (ECA437) contains one N2 locus and one CB4856 locus, and the resistant strain (ECA481) contains two N2 loci. Because this region is in the center of a chromosome where recombination frequency is lower [33], we were unable to generate NILs with a breakpoint to further narrow the QTL. Furthermore, it is possible that multiple small-effect loci are contributing to each of the two QTL, rendering it difficult to identify each causal gene or variant. Regardless, at least two novel loci on chromosome V were identified that influence zinc sensitivity in *C. elegans*.

**Fig 2.**
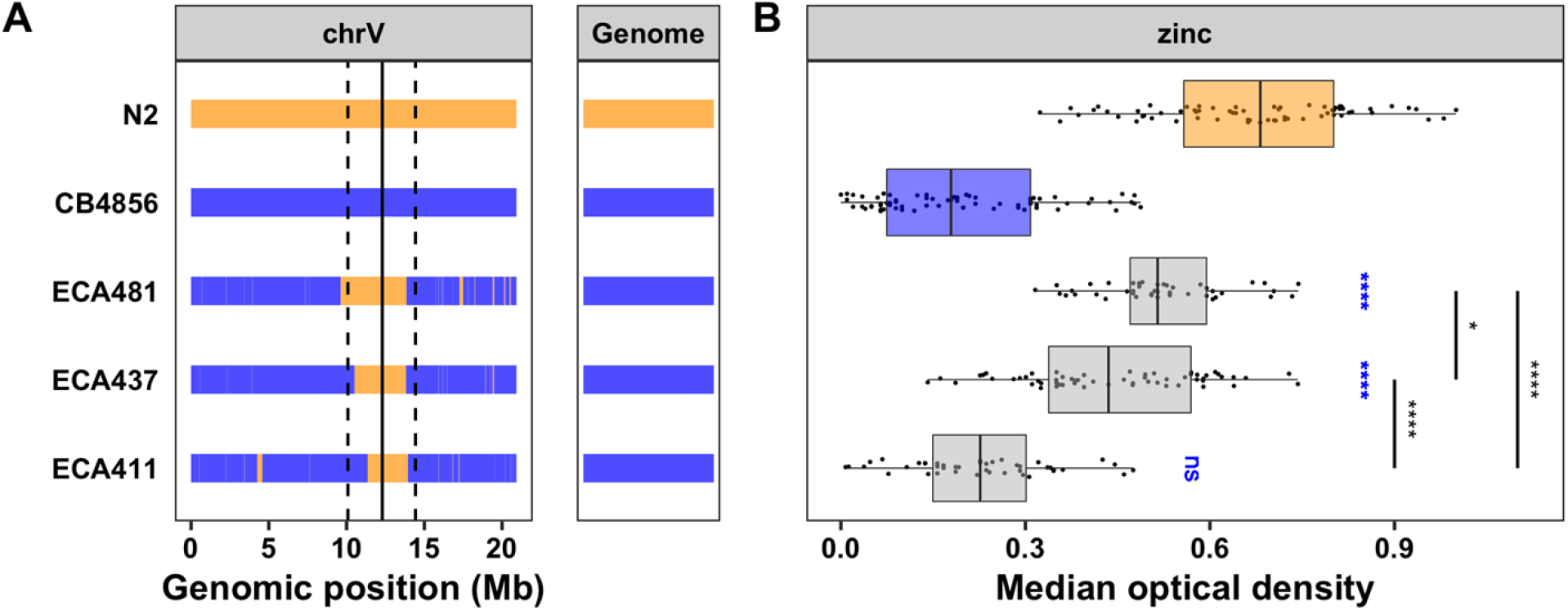
NILs identify multiple QTL on chromosome V. **A)** Strain genotypes are shown as colored rectangles (N2: orange, CB4856: blue) in detail for chromosome V (left) and in general for the rest of the chromosomes (right). The solid vertical line represents the peak marker of the QTL, and the dashed vertical lines represent the confidence interval. **B)** Normalized residual median optical density in zinc (median.EXT, x-axis) is plotted as Tukey box plots against strain (y-axis). Statistical significance of each NIL compared to CB4856 is shown above each strain (ns = non-significant (p-value > 0.05); *, **, ***, and *** = significant (p-value < 0.05, 0.01, 0.001, or 0.0001, respectively).

### Analysis of the chromosome III QTL suggests that a sequestosome-related gene, *sqst-5*, contributes to differences in zinc responses

The QTL on chromosome III accounts for 20% of the phenotypic variance in zinc response across the RIAIL population. In contrast to the previous three QTL, the CB4856 allele is associated with zinc resistance and the N2 allele is associated with zinc sensitivity (**Fig 1C, S3 and S4 Files**). The strain with the N2 allele on chromosome III crossed into a CB4856 genetic background (ECA859) was significantly more sensitive than the CB4856 strain (hyper-sensitive) and the strain with the opposite genotype (ECA838) was significantly more resistant than the N2 strain (hyper-resistant) (**S5 Fig, S9 and S10 Files**). These results demonstrate that this locus contributes to the observed transgressive phenotypes in the RIAILs (**Fig 1A, S3 File**). We also measured animal optical densities in zinc for individuals heterozygous for the chromosome III locus to determine whether the N2 or CB4856 allele confers the dominant phenotype. To analyze heterozygous individuals, we developed a modified high-throughput assay (see Methods, **S6 Fig, S12 File**). Individuals heterozygous for the chromosome III locus in the N2 genetic background (N2xECA838) were hyper-resistant similar to the NIL that is homozygous CB4856 for the chromosome III locus in the N2 genetic background (ECA838) (**Fig 3, S10 and S13 Files**). By contrast, individuals heterozygous for the chromosome III locus in the CB4856 genetic background (CB4856xECA859) were significantly more resistant than their hyper-sensitive NIL counterpart, which is homozygous N2 for the chromosome III locus in the CB4856 genetic background (ECA859). The phenotype of this heterozygous strain was also similar to that of the CB4856 strain. The results of these crosses validate that genetic variation between N2 and CB4856 on the left arm of chromosome III contributes to the nematode zinc response and indicate that the CB4856 allele conferred a dominant phenotype.

**Fig 3.**
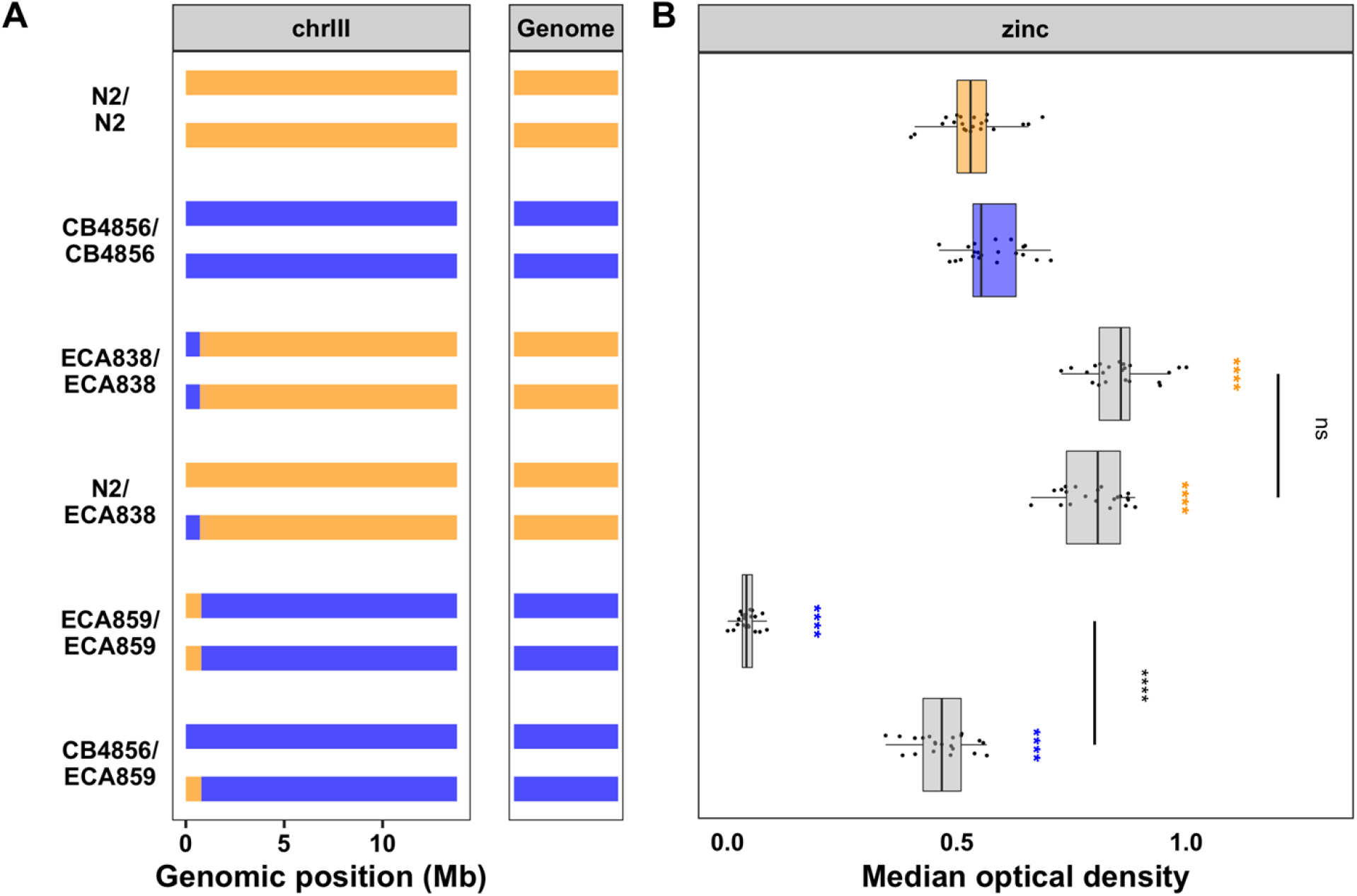
Dominance of chromosome III QTL. **A)** Strain genotypes are shown as colored rectangles (N2: orange, CB4856: blue) in detail for chromosome III (left) and in general for the rest of the chromosomes (right). Each rectangle represents a single copy of chromosome III. **B)** Normalized residual median optical density in zinc (median.EXT, x-axis) is plotted as Tukey box plots against strain (y-axis). The N2 strain, which is normally resistant to zinc, was sick in this experiment. Statistical significance of each strain compared to its parental strain (ECA838/ECA838 and N2/ECA838 to N2 and ECA859/ECA859 and CB4856/ECA859 to CB4856) is shown above each strain and colored by the parent strain it was tested against (ns = non-significant (p-value > 0.05); *, **, ***, and *** = significant (p-value < 0.05, 0.01, 0.001, or 0.0001, respectively).

Because no previously identified zinc-related genes are in this interval, we investigated the composition of the genes in N2 to look for any obvious candidates that might underlie this QTL. We found 119 genes in this interval (**Table 1, S13 File**). A change in phenotype is often observed when either genetic variation causes a change in the amino-acid sequence of the protein (protein-coding variation) or genetic variation causes a change in gene expression. Previously, whole-genome gene expression was measured in a set of 208 RIAILs derived from the N2 and CB4856 strains [50] and expression QTL (eQTL) mapping was performed [50,63]. We used this dataset to find genes with an eQTL that maps to our region of interest. In total, we eliminated 19 genes that had no genetic variation in CB4856 and prioritized 62 genes that had protein-coding variation and/or an eQTL that mapped to this region (**Table 1, S14 File**).

**Table 1:** Genes in QTL intervals for chromosomes II and V

| QTL interval (bp) | No genetic variation<sup>a</sup> | Protein-coding variation and/or eQTL<sup>b</sup> | Other genetic variation<sup>c</sup> | Other eQTL that map to interval<sup>d</sup> | Total |
| --- | --- | --- | --- | --- | --- |
| **III:4,664-597,553** | 19 | 55 | 45 | 7 | **126** |
| **V:9,620,518-10,511,995** | 183 | 49 | 97 | 24 | **353** |
| **V:10,511,995-11,345,444** | 215 | 45 | 115 | 19 | **394** |

<sup>a</sup>Genes within genomic interval with no genetic variation
<sup>b</sup>Genes within genomic interval with protein-coding variation and/or an eQTL that maps to this interval
<sup>c</sup>Genes within genomic interval with non-protein-coding variation and no eQTL that maps to this interval
<sup>d</sup>Genes outside genomic interval with an eQTL that maps to this interval

To narrow our list of genes further, we analyzed the functional descriptions and gene ontology (GO) annotations for all 62 candidate genes. A gene that is predicted to bind zinc and has protein-coding variation or variation in gene expression between N2 and CB4856 would be a high-priority candidate. We identified four genes that are predicted to bind zinc and a fifth gene that is regulated by a zinc finger transcription factor (**S14 File**). Upon further inspection, one of these five genes (*sqst-5*) also had an eQTL that was originally assigned to the nearby pseudogene *ver-2* (**Fig 4A, S7 Fig and S15 File**). RIAILs with the N2 allele at the *sqst-5* locus have significantly higher expression of the gene than those with the CB4856 allele (**Fig 4B and S15 File**). We previously showed that mediation analysis can be a useful tool to link variation in gene expression with drug-response phenotypes [63]. We used the standard high-throughput assay to measure zinc responses for 121 of the 208 RIAILs with gene expression data (**S3 File**) and performed mediation analysis for each of the 17 genes with an eQTL in the region (**S16 File**). The mediation estimate for *sqst-5* was the strongest hit (**Fig 4C**). Together, these results suggest that genetic variation on chromosome III causes a decrease in expression of *sqst-5* that leads to increased zinc resistance.

**Fig 4.**
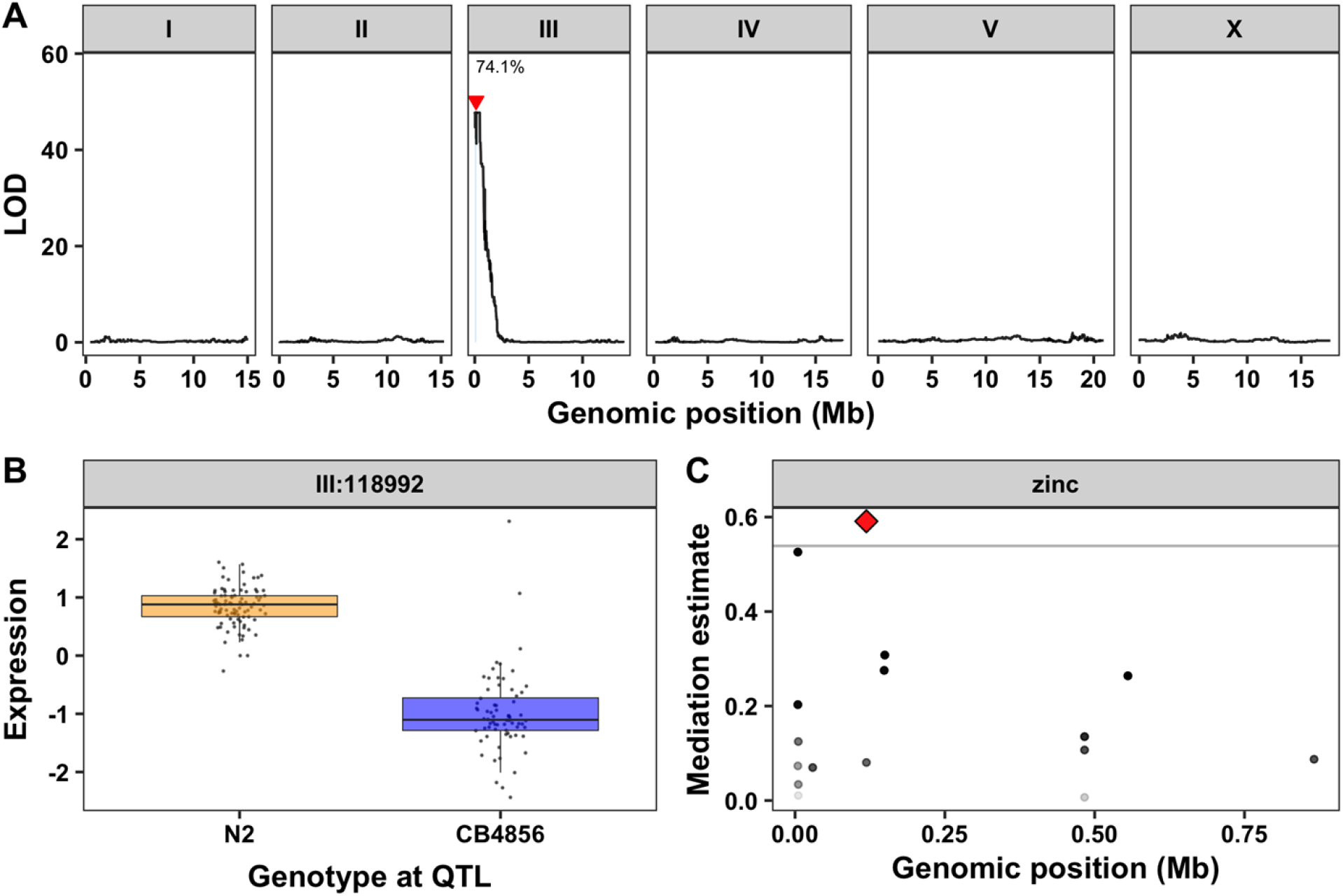
Expression QTL mapping and mediation analysis for *sqst-5*. **A)** Results from the linkage mapping using expression of *sqst-5* as a quantitative trait. Genomic position (x-axis) is plotted against the logarithm of the odds (LOD) score (y-axis) for 13,003 genomic markers. The significant QTL is indicated by a red triangle at the peak marker, and a blue rectangle shows the 95% confidence interval around the peak marker. The percentage of the total variance in the RIAIL population that can be explained by the QTL is shown above the QTL. **B)** The expression of *sqst-5* (y-axis) of RIAILs split by genotype at the marker with the maximum LOD score (x-axis) are plotted as Tukey box plots. Each point corresponds to a unique recombinant strain. Strains with the N2 allele are colored orange, and strains with the CB4856 allele are colored blue. **C)** Mediation estimates calculated as the indirect effect that differences in expression of each gene plays in the overall phenotype (y-axis) are plotted against genomic position of the eQTL (x-axis) on chromosome III for al 17 genes with an eQTL in the drug-response QTL confidence interval. The 95th percentile of the distribution of mediation estimates is represented by the horizontal grey line. The confidence of the estimate increases (p-value decreases) as points become more solid. *sqst-5* is represented by a red diamond.

### Variation in *sqst-5* underlies differences in zinc responses

To test the function of *sqst-5* in the zinc response, we constructed two independently derived strains harboring large deletions of *sqst-5* in both the N2 and CB4856 genetic backgrounds. Because RIAILs with the CB4856 allele (which was associated with higher resistance to zinc) have lower expression of *sqst-5* (**Fig 4B and S15 File**), we expected *sqst-5* deletions in the CB4856 genetic background might cause little or no change in zinc resistance. Alternatively, we expected *sqst-5* deletions in the N2 genetic background might cause increased zinc resistance. Surprisingly, we found that deletions of *sqst-5* had no effect in either background (**S8 Fig, S10 File and S17 Files**). However, the increased sensitivity of the N2 allele in the CB4856 genetic background (ECA859) always had a much larger effect than the increased resistance of the CB4856 allele in the N2 background (ECA838) (**Fig 3, S5 Fig, S9-S10 and S13 Files**). To take advantage of this sensitization, we deleted *sqst-5* in the hyper-sensitive NIL strain that contains the N2 *sqst-5* allele in the CB4856 genetic background (ECA859). We hypothesized that deleting *sqst-5* in the hyper-sensitive NIL would make this strain less sensitive to zinc (more similar to the CB4856 phenotype). As expected, we observed a significant increase in resistance for these deletions compared to the NIL (**S9 Fig, S10 and S18 Files**), indicating a role for *sqst-5* in the *C. elegans* zinc response.

To provide further evidence that natural variation between N2 and CB4856 in *sqst-5* underlies the chromosome III QTL, we measured the optical density of individuals hemizygous for the N2 *sqst-5* allele in the hyper-sensitive NIL genetic background (ECA2517xECA859) in response to zinc. If a loss-of-function allele of *sqst-5* in the CB4856 strain is responsible for the variation in zinc response between N2 and CB4856 (**S8 Fig, S10 and S17 Files**), then this hemizygous strain should show the same sensitivity as both the CB4856 strain and the strain with the homozygous deletion of *sqst-5* in the hyper-sensitive NIL genetic background (ECA2517). We observed that the strain hemizygous for the N2 *sqst-5* allele was indeed more resistant than the hyper-sensitive NIL and similar in sensitivity to the CB4856 strain (**Fig 5, S10 and S19 Files**). This result recapitulated the result of the dominance assay (**Fig 3**), suggesting that a loss-of-function allele of *sqst-5* conferred a dominant resistance phenotype. A dominant phenotype caused by a loss-of-function allele is, most times, caused by haploinsufficiency. Therefore, the zinc-response phenotype driven by the single functional N2 *sqst-5* allele is not sufficient to produce the hyper-sensitive phenotype of the NIL with two functional N2 alleles.

**Fig 5.**
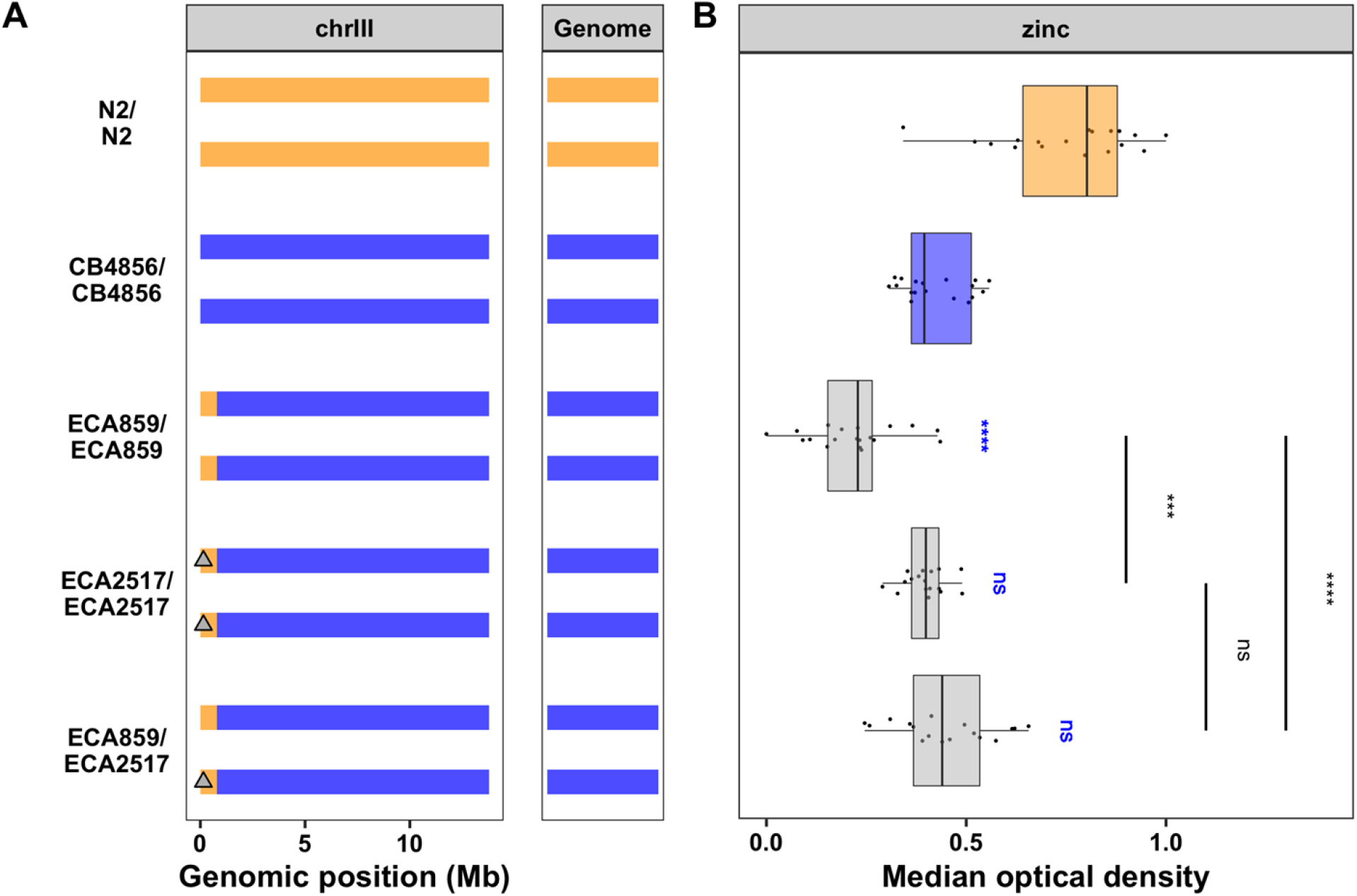
The gene *sqst-5* confers zinc sensitivity. **A)** Strain genotypes are shown as colored rectangles (N2: orange, CB4856: blue) in detail for chromosome III (left) and in general for the rest of the chromosomes (right). Each rectangle represents a single copy of chromosome III, and grey triangles represent *sqst-5* deletions. **B)** Normalized residual median optical density in zinc (median.EXT, x-axis) is plotted as Tukey box plots against strain (y-axis). The N2 strain, which is normally resistant to zinc, was sick in this experiment. Statistical significance of each strain compared to CB4856 is shown above each strain and NIL pairwise significance is shown as a bar above strains (ns = non-significant (p-value > 0.05); *, **, ***, and *** = significant (p-value < 0.05, 0.01, 0.001, or 0.0001, respectively).

### A natural deletion in *sqst-5* is conserved across wild isolates

We next searched for specific genetic variants in *sqst-5* that could lead to a loss-of-function allele in CB4856. We investigated the sequence read alignments of the N2 and CB4856 strains at the *sqst-5* locus using the Variant Browser on CeNDR (elegansvariaton.org) [32] and observed a putative large deletion in the second exon. We confirmed that this deletion is 111 bp (N2 coordinates: chrIII:147,076-147,186 bp) using a whole-genome alignment between the N2 reference genome and the CB4856 genome recently assembled using long-read sequencing [65] (**Fig 6**, **S20 and S21 Files**). We ran gene prediction algorithms on the CB4856 sequence, but no gene was predicted (**S22 File**). The SQST-5 protein in the N2 strain has a single characterized protein domain: a Zinc finger, ZZ-type (Wormbase.org, WS275). The ZZ-type domains are predicted to bind two zinc ions using a repeated conserved motif of Cys-X2-Cys and are also important for protein-protein interactions [66,67]. Interestingly, when we overlaid the location of the ZZ-type domain with the CB4856 alignment, we discovered that the 111 bp deletion spans most of the ZZ-type domain, including the essential Cys-X2-Cys motif (**Fig 6A**). Because this domain is important for binding zinc ions, this result suggests that even if CB4856 expresses low levels of SQST-5, it is unlikely to bind zinc at the same level as strains with a complete ZZ-type domain.

**Fig 6.**
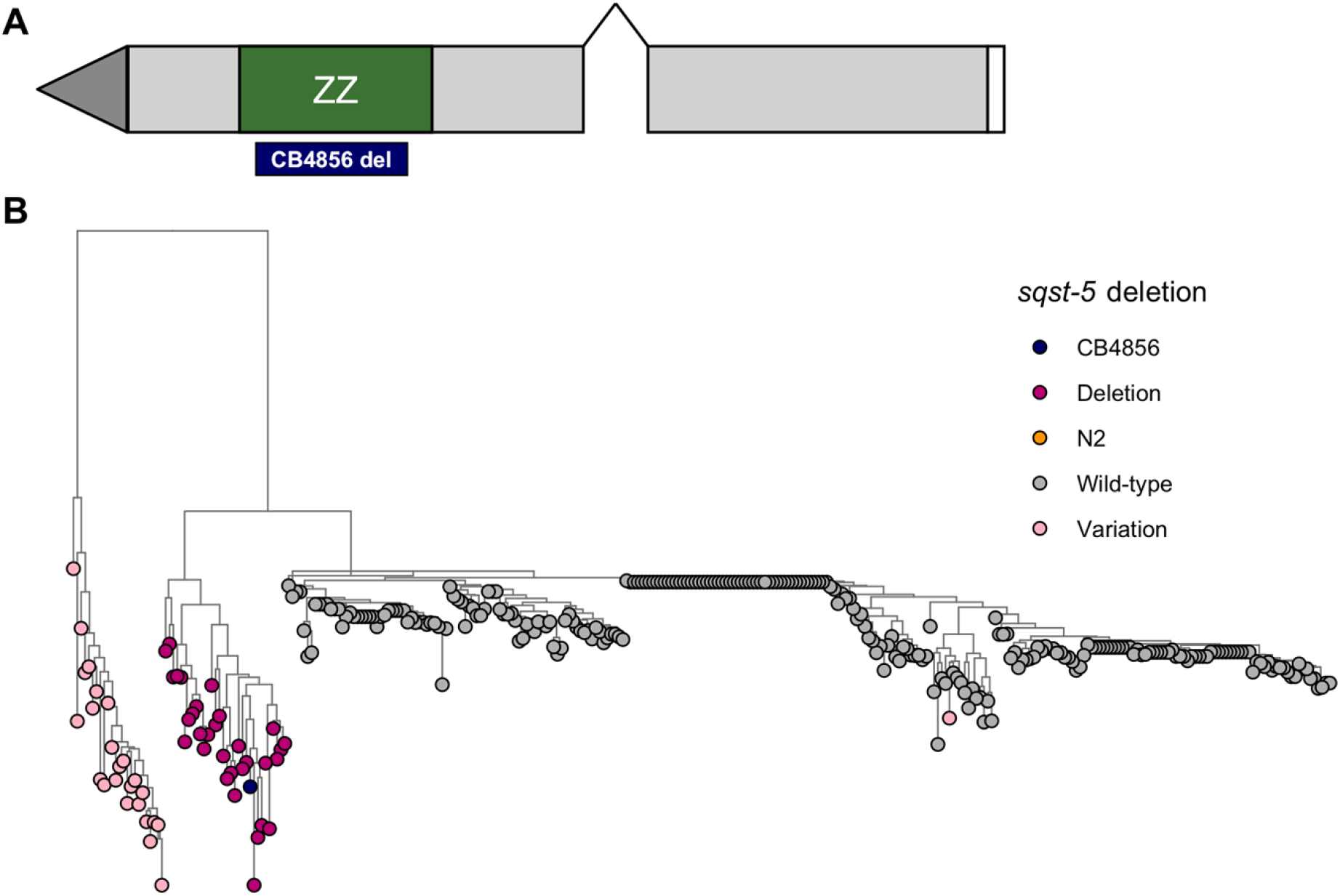
Natural genetic variation in *sqst-5*. **A)** Gene model of *sqst-5*. Grey rectangles represent exons, and connecting black lines represent introns. The ZZ-type domain is indicated by a green rectangle, and the location of the natural 111 bp deletion in CB4856 is indicated below with a blue rectangle. **B)** Neighbor-joining tree indicating genetic relatedness between wild isolates at the *sqst-5* locus. Branch lengths indicate the rate, the tree is midpoint rooted. Tips are colored by the variation haplotype at *sqst-5*: (Wild-type: grey, N2: orange, CB4856: navy, Deletion: magenta, Other putative structural variation: light pink).

We next investigated structural variation across a panel of 328 wild isolates to ask if this deletion is unique to the CB4856 strain or common across many wild strains. We identified 31 additional strains with the same 111 bp deletion as CB4856 by manual inspection using the Variant Browser on CeNDR (**S23 File**). We also identified 25 strains that harbored low sequence identity with the N2 reference genome, indicating that these strains might contain structural variation different from the deletion in the CB4856 strain. We assessed the genetic relatedness of these strains by constructing a neighbor-joining tree for all 328 wild isolates using the single nucleotide variants near and within *sqst-5* (**S24 File**). All 32 strains with the predicted deletion in *sqst-5* cluster together (**Fig 6B**), suggesting these strains inherited this deletion from a common ancestor. These strains were not isolated from a single location, but rather spread geographically across Europe and the Pacific Rim (**S10 Fig and S23 File**). Furthermore, 24 of the 25 strains with other putative structural variation in *sqst-5* also cluster together separately from the strains with the 111 bp deletion (**Fig 6B and S24 File**). This result suggests that this second group of strains also share a common ancestor that harbored variation in *sqst-5*. Additionally, strains with the deletion and strains with the other haplotype are sometimes found in nearby locations (**S10 Fig and S23 File**). Regardless, the 111 bp deletion and putative other structural variants might cause loss of *sqst-5* function.

To test if a loss-of-function allele of *sqst-5* correlates with zinc resistance among our panel of wild isolates, we measured animal development (length, optical density, and normalized optical density) and reproductive ability (brood size) of 81 strains in response to zinc (**S25 File and S25 File**). Including CB4856, we tested nine strains with variation in *sqst-5:* four strains with the 111 bp deletion and five strains with the other putative structural variation. On average, these nine strains were more resistant than the rest of the population (**Fig 7A, S11 Fig and S26 File**), and variation in *sqst-5* explained up to 11.5% (median.EXT; *p-value = 0.0019*) of the total variation in zinc responses among the wild isolates. Genome-wide association mapping identified eight small-effect QTL across the four zinc-response traits (**S11 Fig**). One trait, normalized optical density, had a QTL on the left arm of chromosome III (**Figs 7B-C**). The proximity of this QTL to *sqst-5* suggests that natural variation in *sqst-5* likely also contributes to variation in response to zinc among a panel of wild *C. elegans* strains.

**Fig 7.**
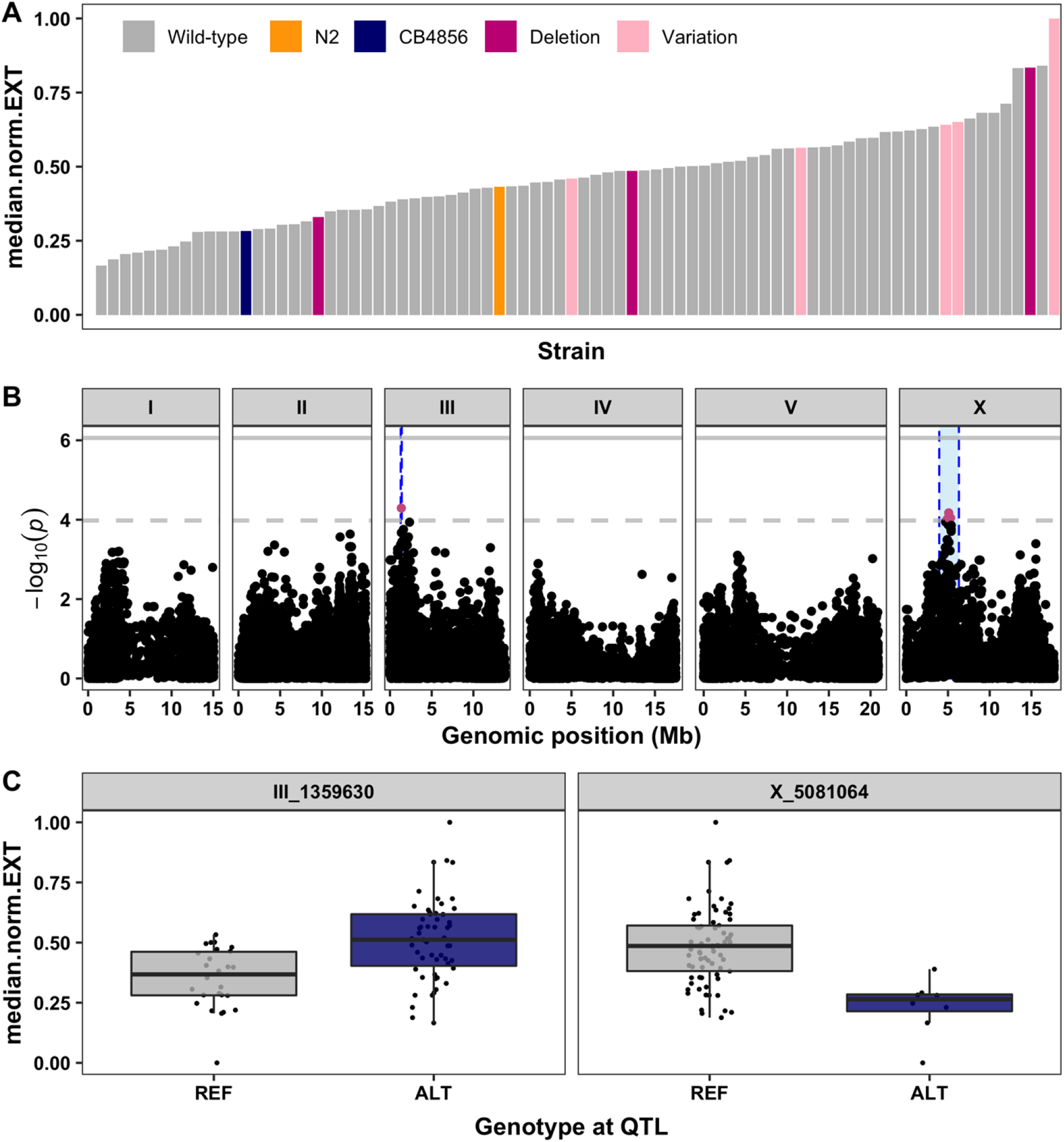
Genome-wide association (GWA) mapping suggests common variation at *sqst-5* is associated with differences in zinc responses among wild isolates. **A)** Normalized residual median normalized optical density (median.norm.EXT, y-axis) of 81 wild isolates (x-axis) in response to zinc supplementation. Strains are colored by the parental strains N2 (orange) and CB4856 (blue) or by the *sqst-5* variation (Deletion: magenta, other variation: light pink) **B)** GWA results are shown. Genomic position (x-axis) is plotted against the *-log10(p)* value (y-axis) for each SNV. SNVs are colored pink if they pass the genome-wide eigen-decomposition significance threshold designated by the dotted grey line. The solid grey line represents the more stringent Bonferroni significance threshold. The genomic regions of interest that pass the significance threshold are highlighted by blue rectangles. **C)** For each QTL, the normalized residual median normalized optical density (median.norm.EXT, y-axis) of strains split by genotype at the peak marker (x-axis) are plotted as Tukey box plots. Each point corresponds to a wild isolate strain. Strains with the N2 reference allele are colored grey, and strains with an alternative allele are colored navy.

## Discussion

We used linkage mapping to identify four QTL in response to high levels of exogenous zinc. We validated the two QTL with the largest effects on chromosomes III and V using near-isogenic lines. The QTL on chromosome V was further dissected into at least two additive loci that became difficult to narrow further. Mediation analysis was performed for the genes with expression variation that overlaps with the QTL on chromosome III, and a single gene (*sqst-5*) was identified whose variation in expression between N2 and CB4856 is correlated with differences in responses to zinc. The CB4856 strain harbors a natural deletion of this gene. We deleted the N2 version of *sqst-5* using CRISPR-Cas9 genome editing and showed that strains without *sqst-5* were significantly more resistant to exogenous zinc than strains with a functional copy of *sqst-5*, suggesting a new role for *sqst-5* in zinc homeostasis. In addition to CB4856, several other wild isolates were found to have a putative independent structural variant in *sqst-5* that is loosely correlated with resistance to zinc. These strains cluster genetically into two distinct groups, suggesting that functional variation in *sqst-5* has emerged multiple times. These results demonstrate the power of leveraging natural genetic variation to identify novel genes in a toxin-response pathway and suggest mechanisms for how high exogenous zinc can be mitigated in natural environments.

### A complex genetic architecture underlies differences in zinc responses

The identification of multiple QTL in response to excess zinc is not surprising, as zinc is an essential trace element and up to 8% of *C. elegans* genes have been predicted to encode proteins that bind zinc [22]. In particular, 28 genes encode putative zinc transporters and an additional 15 genes have been identified via mutagenesis screens in N2 that promote either zinc resistance or sensitivity [8,18]. We identified two QTL that contain at least one gene that was previously found to be involved in the nematode zinc response and an additional two QTL that do not contain any genes previously found to affect zinc responses (**S6 File**). Several of the genes with previously described roles in the zinc response are located on chromosome X. However, only three of these genes (*hke-4.2, hizr-1*, and *elt-2*) have protein-coding variation in CB4856. The results of the linkage mapping experiment (**Fig 1B**) identified a broad peak on chromosome X, but we were unsuccessful validating this QTL using NILs (**S5 Fig**) likely because the QTL could contain multiple small-effect loci that could each act in opposite directions. By contrast, we were able to validate that genetic variation on chromosome V contributes to the nematode zinc responses (**Fig 2**). However, as more NILs were tested with smaller introgressions, we observed a fractionation of the QTL into at least two small-effect loci. Several previous studies that aimed to deeply validate a single QTL have instead identified many tightly linked antagonistic QTL underlying the major QTL [49,68–75]. Unfortunately, as each QTL fractionates into several QTL, the individual effect sizes become smaller and our ability to accurately interpret signal from noise becomes more difficult. This polygenic nature of complex traits is a major roadblock in going from QTL to QTG [76–78].

Although we were unable to identify specific genes on chromosome V that contribute to the nematode zinc response, we were able to narrow the QTL interval from 4.3 Mb (chrV:10,084,029-14,428,285) to 1.2 Mb (chrV:10,084,029-11,345,444) containing at least two loci that underlie differential responses to zinc. Of the 573 genes of interest (**Table 1**), we identified two high-priority candidate genes (*zhit-3* and *H27A22.1*) that are predicted to bind zinc and have protein-coding variation in the CB4856 strain. The gene *zhit-3* encodes a protein that is an ortholog of the human protein ZNHIT6 and contains a zinc finger HIT-type domain (Wormbase.org, WS275). The gene *H27A22.1* encodes a protein that is an ortholog of the human protein QPCTL with glutaminyl-peptide cyclotransferase (Wormbase.org, WS275). It is possible that genetic variation in one or both of these genes underlies the QTL on chromosome V. However, future studies are needed to confirm the role of these genes in the nematode zinc response.

Although zinc is the most biologically relevant heavy metal, other divalent cations with similar chemistries also play important roles in biological systems (copper, nickel, and iron) or are highly toxic (cadmium) [1,2,8]. We previously performed linkage mapping for three of these heavy metals and found that QTL for zinc, copper, and nickel overlap on the right arm of chromosome IV (**S12 Fig and S27 File**) [43,75]. The overlap of this QTL with other heavy metal QTL suggest that perhaps the molecular mechanisms underlying these QTL are not specific to zinc. Furthermore, this QTL is in regions previously defined as a QTL hotspot where a single pleiotropic gene might control several toxin responses or several independent yet tightly linked genes might control different traits [43]. Regardless, the high-effect QTL on chromosome III seems to be unique to the zinc response, as none of the other metals have a QTL on chromosome III (**S12 Fig**).

### Common genetic variation underlies differential responses to exogenous zinc

We discovered 31 additional wild strains with the same 111 bp deletion in *sqst-5* as found in the CB4856 strain and another 25 strains that show evidence of different structural variation. Long-read sequencing and local genome assembly of strains with this alternative variation are needed to fully define these haplotypes. Although these strains were isolated globally (**S10 Fig**), phylogenetic analysis suggests that these strains comprise two common classes of variation at the *sqst-5* locus (**Fig 6B**). Strains from these two classes of variation are sometimes found in close geographical proximity, indicating a possibility for convergent evolution in zinc resistance, perhaps in geographic regions with high levels of zinc in the environment. We require further investigation of metal contents in niches that contain *C. elegans* to connect environmental zinc levels to natural genetic variation.

We measured zinc responses for a subset of wild strains and found that variation in *sqst-5* could explain 11.5% of the total variation in response to zinc among the panel of wild isolates. Interestingly, N2 and CB4856 were among the more sensitive strains tested (**Fig 7 and S11 Fig**), suggesting the existence of several loci not found within the CB4856 strain that influence zinc responses. GWA mapping discovered several small-effect loci across the genome that were associated with zinc resistance or sensitivity (**S11 Fig**). In particular, the QTL on chromosome III (nearby the *sqst-5* locus) and chromosome X overlapped with QTL discovered using linkage mapping. Alternatively, a QTL on the right arm of chromosome III provides evidence of common genetic variation not present in the CB4856 strain that plays a role in the nematode zinc response. Because none of the known zinc-related genes are in this interval, this QTL might represent another novel gene that contributes to zinc resistance or sensitivity (**S6 File**). Our power to detect QTL would only improve with the phenotyping of more strains in the presence of high exogenous zinc.

### SQST-5 might function to negatively regulate other sequestosome-related genes

We show that strains with a functional copy of *sqst-5* are more sensitive to zinc than strains with a large deletion of the gene, indicating that *sqst-5* negatively regulates the zinc response. Using BLASTp (Wormbase.org, WS275), we searched for paralogs and identified five other members of the sequestosome-related family (*sqst-1, sqst-2, sqst-3, sqst-4*, and *C06G3.6*) each containing a ZZ-type domain, like *sqst-5*. Two of these genes have been previously implicated in the nematode stress response. The gene *sqst-1* is upregulated in response to hormetic heat shock [79]. Both SQST-1 and the human ortholog, SQSTM1/p62, have been shown to bind to and target ubiquitinated proteins to an organelle (sequestosome) for subsequent degradation by autophagy [79,80]. The ZZ-domain, particularly the zinc-coordinating Cys-X2-Cys residues, has been shown to be essential for this process [81,82]. Additionally, *sqst-3* is expressed in response to exogenous cadmium [83], suggesting that the sequestosome-related family might be involved in divalent cation metal stress responses. If we connect these disparate results, then these genes could protect cells against zinc toxicity using sequestration and fusion with lysosomes. Alternatively, divalent cation metal stress could cause disruption of proteostasis and upregulation of sequestosome genes indirectly related to the specific metal stress. The role of this gene family in stress response has not been characterized using loss-of-function genetics, so we do not know whether the family is protective in response to exogenous stressors like zinc.

If the sequestosome-related genes do function to protect cells from high exogenous zinc, we would expect that loss-of-function of these genes should cause increased zinc sensitivity. However, loss-of-function of *sqst-5* caused increased zinc resistance, indicating that *sqst-5* might have a different function than other sequestosome-related genes. Although the exact function of SQST-5 is unknown, it is predicted to have protein kinase C and K63-linked polyubiquitin modification-dependent protein binding activity (Wormbase.org, WS275). From previous studies, we know that the ZZ-type domain is important for protein-protein interactions [66], and we discovered that the natural 111 bp deletion in the CB4856 strain causes a loss of this zinc-binding domain. It is possible that SQST-5 could function as an inhibitor of mechanisms that mitigate exogenous zinc, potentially by binding to other sequestesome-related proteins and inhibiting their activities. When SQST-5 is removed, the protein partner is no longer inhibited and is available to bind or capture the excess zinc in the environment, thus reducing the toxicity induced by high exogenous zinc. Biological processes that are finely balanced in homeostasis often contain both positive and negative regulators. Functional studies to directly test the role of *sqst-5* in the autophagy pathway, both in control conditions and in the presence of exogenous zinc, are necessary to provide insights into its function as a negative regulator of zinc homeostasis. Likewise, zinc responses of animals with targeted deletions of the other sequestosome-related genes are needed to fully define the roles for these genes in the *C. elegans* zinc response pathway. Overall, this study leverages natural genetic variation to discover a novel gene that sensitizes nematodes to exogenous zinc, potentially by creating a negative feedback loop to regulate other sequestosome-related genes.

## Materials and methods

### Strains

Animals were grown at 20°C on modified nematode growth media (NGMA) containing 1% agar and 0.7% agarose to prevent burrowing and fed OP50 [44]. The two parental strains, the canonical laboratory strain, N2, and the wild isolate from Hawaii, CB4856, were used to generate all recombinant lines. 253 recombinant inbred advanced intercross lines (RIAILs) generated previously [34] (set 2 RIAILs) were used for zinc phenotyping and QTL mapping. A second set of 121 RIAILs generated previously [33] (set 1 RIAILs) were phenotyped for mediation analysis. All strains are listed in the Supplementary Material and are available upon request or from the *C. elegans* Natural Diversity Resource [32].

### Standard high-throughput fitness assay

For dose responses and RIAIL phenotyping, we used a high-throughput fitness assay (HTA) described previously [34]. In summary, populations of each strain were passaged and amplified on NGMA plates for four generations without starvation. In the fifth generation, gravid adults were bleach-synchronized and 25-50 embryos from each strain were aliquoted into 96-well microtiter plates at a final volume of 50 μL K medium [85]. The following day, arrested L1s were fed HB101 bacterial lysate (Pennsylvania State University Shared Fermentation Facility, State College, PA; [86]) at a final concentration of 5 mg/mL in K medium and were grown to the L4 larval stage for 48 hours at 20°C with constant shaking. Three L4 larvae were sorted into new 96-well microtiter plates containing 10 mg/mL HB101 bacterial lysate, 50 μM kanamycin, and either 1% water or zinc sulfate dissolved in 1% water using a large-particle flow cytometer (COPAS BIOSORT, Union Biometrica; Holliston, MA). Sorted animals were grown for 96 hours at 20°C with constant shaking. The next generation of animals and the parents were treated with sodium azide (50 mM in 1X M9) to straighten their bodies for more accurate measurements. Animal length (TOF) and optical density (EXT) were quantified for every animal in each well using the COPAS BIOSORT and the medians of each well population (median.TOF and median.EXT) were used to estimate these traits. Animal length and optical density are both measures of nematode development; animals get longer and more optically dense (thicker and denser body composition) as they develop [34]. However, the COPAS BIOSORT measures optical density as a function of length. Because these two traits are highly correlated, we also generated a fourth trait (median.norm.EXT) which normalizes the optical density by length (EXT/TOF) in order to provide a means to compare optical densities regardless of animal lengths. Finally, brood size (norm.n) was calculated as the total number of animals in the well normalized by the number of parents originally sorted and provides an estimate of nematode reproductive fitness [34].

### Calculation of zinc-response traits

Phenotypic measurements collected by the BIOSORT were processed and analyzed using the R package *easysorter* [87] as described previously [35]. Briefly, raw data from the BIOSORT was read into R using the *read_data* function and contaminated wells were removed using the *remove_contamination* function. The *sumplate* function was used to calculate summary statistics per well and four main traits were output: median.TOF (animal length), median.EXT (animal optical density), median.norm.EXT (animal optical density normalized by animal length), and norm.n (brood size). When trait measurements were collected across multiple assay experiments, the *regress(assay = T*) function was used to fit the linear model (*phenotype ~ assay*) to account for variation among assays. Outliers were pruned using *prune_outliers* to remove wells beyond two standard deviations of the mean for highly replicated assays. Alternatively, for assays with low replication (dose response and RIAIL phenotyping), *bamf_prune* was used to remove wells beyond two times the IQR plus the 75th quartile or two times the IQR minus the 25th quartile, unless at least 5% of the strains lie outside this range. Finally, zinc-specific effects were calculated using the *regress(assay = FALSE*) function, which subtracts the mean water (control) value from each zinc replicate for each strain using a linear model (*drug_phenotype ~ control_phenotype)*. The residual phenotypic values were used as the zinc-response phenotype for all downstream analyses. In this way, we addressed only the differences among strains that were caused by treatment with zinc and ignored minor phenotypic variation among strains in the control condition. Pairwise tests for statistically significant differences in the zinc response between strains were performed using the *TukeyHSD* function [88] on an ANOVA model with the formula (*phenotype ~ strain)*. For plotting purposes, these residual values were normalized from zero to one with zero being the well with the smallest value and one the well with the largest value.

### Zinc dose response

Four genetically divergent strains (N2, CB4856, JU258, and DL238) were treated with increasing concentrations of zinc using the standard high-throughput assay described above. A concentration of 500 μM zinc sulfate heptahydrate (Sigma #221376-100G) in water was selected for the linkage mapping experiments. This concentration provided a reproducible zinc-specific effect and maximizes between-strain variation and minimizes within-strain variation across the four traits. Broad-sense heritability was calculated from the dose response phenotypes using the *lmer* function in the *lme4* R package [89] with the formula *phenotype ~ 1 + (1 | strain*) for each dose.

### Linkage mapping

253 RIAILs (set 2 RIAILs) were phenotyped in both zinc and water using the standard high-throughput assay described above. Linkage mapping was performed for all four zinc-response traits using the R package *linkagemapping* (https://github.com/AndersenLab/linkagemapping) as described previously [35]. The cross object derived from the whole-genome sequencing of the RIAILs containing 13,003 SNPs was loaded using the function *load_cross_obj(“N2xCB4856cross_full’)*. The RIAIL phenotypes were merged into the cross object using the *merge_pheno* function with the argument *set = 2*. A forward search (*fsearch* function) adapted from the *R/qtl* package [90] was used to calculate the logarithm of the odds (LOD) scores for each genetic marker and each trait as *-n(ln(1-R^2^)/2ln(10))* where R is the Pearson correlation coefficient between the RIAIL genotypes at the marker and trait phenotypes [91]. A 5% genome-wide error rate was calculated by permuting the RIAIL phenotypes 1000 times. The marker with the highest LOD score above the significance threshold was selected as the QTL then integrated into the model as a cofactor and mapping was repeated iteratively until no further significant QTL were identified. Finally, the *annotate_lods* function was used to calculate the effect size of each QTL and determine 95% confidence intervals defined by a 1.5 LOD drop from the peak marker using the argument *cutoff = chromosomal*. In the same manner, linkage mapping was performed for three other divalent cation metals: 250 μM copper in water, 100 μM cadmium in water [43], and 350 μM nickel in water [75].

### Two-dimensional genome scan

A two-dimensional genome scan to identify interacting loci was performed for animal optical density (median.EXT) in zinc using the *scantwo* function from the *qtl* package [90] as described previously [35,43]. Each pairwise combination of loci was tested for correlations with trait variation in the RIAILs. A summary of the maximum interactive LOD score for each chromosome pair can be output using the *summary* function. Significant interactions were identified by permuting the phenotype data 1000 times and determining the 5% genome-wide error rate. The significant interaction threshold for the zinc-response variation scantwo was 4.09.

### Construction of near-isogenic lines (NILs)

NILs were generated as previously described [35,41–43] by either backcrossing a selected RIAIL for six generations or *de novo* by crossing the parental strains N2 and CB4856 to create a heterozygous individual that is then backcrossed for six generations. PCR amplification of insertion-deletion (indel) variants between N2 and CB4856 were used to track the genomic interval. Smaller NILs to further break up the interval were created by backcrossing a NIL for one generation to create a heterozygous F_1_ individual. The F_1_ individuals were selfed, and the F_2_ population was scored for recombination events. NILs were whole-genome sequenced to verify introgressions and the absence of other introgressed regions [35,43]. Reagents used to generate NILs and a summary of each introgression can be found in the **Supplemental Material**.

### Development of R Shiny application to visualize NIL phenotypes

An R shiny web app (version 1.4.0.2) was developed to visualize the results from the high-throughput assays and can be found here: https://katiesevans9.shinyapps.io/QTL_NIL/. To begin analysis, the user can find all data controls in a panel on the left-hand side of the screen. A test dataset is provided (user should check “Use sample data” checkbox) or the user can upload one of the supplementary files from the manuscript after downloading to their local computer. The user should select “water” as the control condition and “zinc” as the drug condition. In some cases, the user will need to choose a specific assay to view if multiple options are available. The user also has the option to view one of four drug-response traits (median.EXT, median.TOF, norm.n, and median.norm.EXT). Finally, the user should choose a chromosome to view, generally the chromosome which contains the highlighted QTL for that particular assay.

Along the top of the main panel, the user can navigate several tabs including “Control”, “Condition”, “Regressed”, “NIL Genotypes”, and “Help!”. The “Control”, “Condition”, and “Regressed” tabs each show the NIL genotypes along the selected chromosome (left) and the NIL phenotypes for the selected trait (right) in the water condition (control), raw zinc condition (condition), or regressed zinc condition (regressed). The regressed data are shown in the manuscript. The user can hover their mouse above the NIL genotype or phenotype plots to see more information or zoom in on a specific area of the plot. Below this plot is an interactive datatable containing the pairwise strain comparisons for this condition and trait. The “NIL Genotypes” tab shows a plot of the NIL genotypes across all chromosomes, not just the chromosome selected by the user (top) and an interactive datatable with the genotypes of each strain across all chromosomes (bottom). The final tab, “Help!” provides the user with the instructions detailed here to help them use the Shiny App.

In addition to these basic controls, the user also has access to several advanced features. The user can choose a subset of strains to plot by checking the box labeled “Show a subset of strains?” and unchecking the boxes next to strains the user wishes to omit. Additionally, the user can plot the location of one or more QTL as a vertical line on the NIL genotype plot by checking the “Show QTL?” box. The user then chooses how many QTL to show and uses the appropriate slider input below to designate the genomic positions of each QTL. Finally, if the “Show genotype?” box is checked, the genotype of each strain at each QTL position will be shown on the phenotype plot as an orange “N” representing N2 and a blue “C” representing CB4856.

### Mediation analysis

121 RIAILs (set 1 RIAILs) were phenotyped in both zinc and water using the standard high-throughput assay described above. Microarray expression for 14,107 probes were previously collected from the set 1 RIAILs [50], filtered [44], and mapped using linkage mapping with 13,003 SNPs [63]. Mediation scores were calculated with bootstrapping using the *mediation* R package [92] as previously described [63] for each of the 19 probes (including *ver-2/sqst-5*, A_12_P104472) with an eQTL on the left arm of chromosome III. Briefly, a mediator model (*expression ~ genotype*) and an outcome model (*phenotype ~ expression + genotype*) were used to calculate the proportion of the QTL effect that can be explained by variation in gene expression. All expression and eQTL data can be found at https://github.com/AndersenLab/scb1_mediation_manuscript.

### Generation of deletion strains

Deletion alleles for *sqst-5* and *ver-2* were generated as previously described using CRISPR-Cas9 genome editing [35,93]. Briefly, 5’ and 3’ guide RNAs were designed with the highest possible on-target and off-target scores [94] and ordered from Synthego (Redwood City, CA). The following CRISPR injection mix was assembled and incubated for an hour before injection: 1 μM *dpy-10* sgRNA, 5 μM of each sgRNA for the gene of interest, 0.5 μM of a single-stranded oligodeoxynucleotide template for homology-directed repair of *dpy-10* (IDT, Skokie, IL), and 5 μM Cas9 protein (Q3B Berkeley, Berkeley, CA) in water. Young adults were mounted onto agar injection pads, injected in either the anterior or posterior arm of the gonad, and allowed to recover on 6 cm plates. After 12 hours, survivors were transferred to individual 6 cm plates and allowed to lay embryos. Two days later, the F1 progeny were screened and individuals with Rol or Dpy phenotypes were selected and transferred clonally to new 6 cm plates. After 48 hours, the F1 individuals were genotyped by PCR flanking the desired deletions. Individuals with heterozygous or homozygous deletions were propagated and genotyped for at least two additional generations to ensure homozygosity and to cross out the Rol mutation. Deletion amplicons were Sanger sequenced to identify the exact location of the deletion. This information as well as all reagents used to generate deletion alleles are detailed in the **Supplemental Material**.

### Modified high-throughput fitness assay

Dominance and validation of candidate genes were tested using a modified version of the standard high-throughput assay detailed above as previously described [35,63]. For candidate gene testing, strains were propagated for two generations, bleach-synchronized in six independent replicates, and titered at a concentration of 25-50 embryos per well of a 96-well microtiter plate. For dominance and hemizygosity assays, strains (males and hermaphrodites) were propagated and amplified for two generations. For each cross, 30 hermaphrodites and 60 males were placed onto each of four 6 cm plates and allowed to mate for 48 hours. Mated hermaphrodites were transferred to a clean 6 cm plate and allowed to lay embryos for eight hours. After the egg-laying period, adults were manually removed and embryos were collected by vigorous washing with 1X M9. Embryos were resuspended in K medium and titered to a concentration of 25 embryos per well of a 96-well microtiter plate. For both assays, arrested L1s were fed HB101 bacterial lysate the following day at a final concentration of 5 mg/mL with either water or zinc. After 48 hours of growth at 20°C with constant shaking, nematodes were treated with sodium azide (5 mM in water) prior to analysis of animal length and optical density using the COPAS BIOSORT. Because only one generation of growth was observed, brood size was not calculated. Lower drug concentrations were needed to see the previous effect because of the modified timing of the drug delivery. A concentration of 250 μM zinc in water was used for these experiments.

### Local alignment of the *sqst-5* region using long-read sequence data

To confirm the putative *sqst-5* deletion in CB4856, we aligned the long-read assembly for CB4856 [65] to the N2 reference genome using NUCmer (version v3.1) [95]. Using this alignment, we identified the coordinates of *sqst-5* in CB4856 and extracted the sequence using BEDtools (version v2.29.2) [96]. We aligned the unspliced N2 *sqst-5* transcript sequence (WormBase WS273) and the N2 SQST-5 protein sequence to this extracted CB4856 sequence using Clustal Omega [97] and GeneWise [98], respectively. Gene prediction in CB4856 was run using Augustus [99]. We visually inspected both alignments to identify the length of the deletion in CB4856 and identify the effect of the deletion of the CB4856 SQST-5 protein sequence.

### Assessment of strain relatedness through neighbor-joining tree

Variant data for dendrogram comparisons were assembled by constructing a FASTA file with the genome-wide variant positions across all strains and subsetting to keep only variants near *sqst-5* (III:145917-148620). Genotype data were acquired from the latest VCF release (release 20180517) from CeNDR. Multiple sequence comparison by log-expectation (MUSCLE, version v3.8.31) [100] was used to generate neighbor-joining trees. A second neighbor-joining tree was constructed with all the variants within the QTL confidence interval for comparison (III:4664-597553). Both trees were identical.

### Genome wide association mapping

Eighty-one wild isolates were phenotyped in both zinc and water using the standard high-throughput assay described above. Genome-wide association (GWA) mappings were performed for all four traits using the R package *cegwas2* (https://github.com/AndersenLab/cegwas2-nf) as described previously [41]. Genotype data were acquired from the latest VCF release (release 20180517) from CeNDR. We used BCFtools [101] to filter variants with missing genotypes and variants below a 5% minor allele frequency. We used PLINK v1.9 [102,103] to LD-prune genotypes. The additive kinship matrix was generated from the 64,046 markers using the *A.mat* function in the *rrBLUP* R package [104]. Because these markers have high LD, we performed eigen decomposition of the correlation matrix of the genotype matrix to identify 477 independent tests [41]. We used the *GWAS* function from the *rrBLUP* package to perform genome-wide mapping. Significance was determined in two ways: a strict Bonferroni threshold and a more lenient eigenvalue threshold set by the number of independent tests in the genotype matrix. Confidence intervals were defined as +/− 150 SNVs from the rightmost and leftmost markers that passed the significance threshold.

### Statistical analysis

All statistical tests of phenotypic differences between strains were performed using the *TukeyHSD* function [88] on an ANOVA model with the formula (*phenotype ~ strain)*. The *p-values* for individual pairwise strain comparisons were adjusted for multiple comparisons (Bonferroni). The datasets and code for generating figures can be found at https://github.com/AndersenLab/zinc_manuscript.

## Acknowledgements

We would like to thank Bryn Gaertner, Samuel Rosenberg, and Tyler Shimko, for assistance with mapping zinc sensitivities and members of the Andersen Lab for helpful comments on the manuscript. Additionally, we would like to thank WormBase and the *C. elegans* Natural Diversity Resource (CeNDR) for data critical for our analysis.

## Supporting information captions

**S1 Fig. Dose response with four divergent wild isolates.** Results from the zinc dose response HTA for brood size (norm.n), animal length (median.TOF), animal optical density (median.EXT), and normalized optical density (median.norm.EXT). For each trait, drug concentration (μM) (x-axis) is plotted against phenotype subtracted from control (y-axis), colored by strain (CB4856: blue, DL238: green, JU258: purple, N2: orange). A red asterisk indicates the dose selected for linkage mapping analysis.

**S2 Fig. Linkage mapping identifies 12 QTL across three traits in response to high zinc. A)** Normalized residual phenotype (y-axis) of 253 RIAILs (x-axis) in response to zinc supplementation. The parental strains are colored: N2, orange; CB4856, blue. **B)** Linkage mapping results are shown. Genomic position (x-axis) is plotted against the logarithm of the odds (LOD) score (y-axis) for 13,003 genomic markers. Each significant QTL is indicated by a red triangle at the peak marker, and a blue rectangle shows the 95% confidence interval around the peak marker. The percentage of the total variance in the RIAIL population that can be explained by each QTL is shown above the QTL. **C)** For each QTL, the normalized residual phenotype (y-axis) of RIAILs split by genotype at the marker with the maximum LOD score (x-axis) are plotted as Tukey box plots. Each point corresponds to a unique recombinant strain. Strains with the N2 allele are colored orange and strains with the CB4856 allele are colored blue.

**S3 Fig. Two dimensional genome scan for median optical density (median.EXT) in zinc.** Log of the odds (LOD) scores are shown for each pairwise combination of loci, split by chromosome. The upper-left triangle contains the epistasis LOD scores and the lower-right triangle contains the LOD scores for the full model. LOD scores are colored, increasing from purple to green to yellow. The LOD scores for the epistasis model are shown on the left of the color scale and the LOD scores for the full model are shown on the right.

**S4 Fig. Reaction norm shows additive QTL effects between chromosome III and V.** Normalized residual median optical density in zinc (median.EXT, y-axis) of RIAILs split by genotype at the chromosome III QTL (x-axis) are plotted as the mean of the population +/− the standard deviation, colored by the genotype at the chromosome V QTL.

**S5 Fig. Validating QTL using near-isogenic lines (NILs). A)** Strain genotypes are shown as colored rectangles (N2: orange, CB4856: blue) in detail for each chromosome (left) and in general for the rest of the chromosomes (right). The solid vertical line represents the peak marker of the QTL and the dashed vertical lines represent the confidence interval. **B)** Normalized residual median optical density in zinc (median.EXT, x-axis) is plotted as Tukey box plots against strain (y-axis). The parental strains N2 and CB4856 are colored orange and blue, respectively. NILs are colored grey. Statistical significance of each strain compared to its parental strain (ECA838, ECA240, ECA232, ECA931, and ECA929 to N2 and ECA859, ECA241, ECA230, and ECA828 to CB4856) is shown above each strain and colored by the parent strain it was tested against (ns = non-significant (p-value > 0.05); *, **, ***, and *** = significant (p-value < 0.05, 0.01, 0.001, or 0.0001, respectively).

**S6 Fig. Dose response for modified HTA.** Results from the zinc dose response with the modified HTA for median optical density (median.EXT). Drug concentration (μM) (x-axis) is plotted against phenotype subtracted from control (y-axis), colored by strain (CB4856: blue, N2: orange). A red asterisk indicates the dose selected for further analysis.

**S7 Fig. The gene *sqst-5*, not *ver-2*, has an eQTL.** Original gene model for *ver-2* is shown with colored boxes representing exons connected by lines representing introns. Exons are colored blue for the new gene model for *ver-2* and purple for *sqst-5*. The black rectangles below represent approximate locations of CRISPR-mediated deletions of *ver-2* or *sqst-5*. The location of the microarray probe is designated as a red rectangle below the plot.

**S8 Fig. Testing the role of *sqst-5* in the zinc response. A)** Strain genotypes are shown as colored rectangles (N2: orange, CB4856: blue) in detail for chromosome III (left) and in general for the rest of the chromosomes (right). The dashed vertical line represents the location of *sqst-5* and grey triangles represent *sqst-5* deletions. **B)** Normalized residual median optical density in zinc (median.EXT, x-axis) is plotted as Tukey box plots against strain (y-axis). Statistical significance of each strain compared to its parental strain (ECA838, ECA1377, and ECA1378 to N2 and ECA859, ECA1379, and ECA1380 to CB4856) is shown above each strain and colored by the parent strain it was tested against (ns = non-significant (p-value > 0.05); *, **, ***, and *** = significant (p-value < 0.05, 0.01, 0.001, or 0.0001, respectively).

**S9 Fig. Isolating the effect of *sqst-5* in the zinc response. A)** Strain genotypes are shown as colored rectangles (N2: orange, CB4856: blue) in detail for chromosome III (left) and in general for the rest of the chromosomes (right). The dashed vertical line represents the location of *sqst-5* and grey triangles represent *sqst-5* deletions. **B)** Normalized residual median optical density in zinc (median.EXT, x-axis) is plotted as Tukey box plots against strain (y-axis). The N2 strain, which is usually resistant to zinc, was sick in this experiment. Statistical significance of each strain compared to ECA859 is shown above each strain (ns = non-significant (p-value > 0.05); *, **, ***, and *** = significant (p-value < 0.05, 0.01, 0.001, or 0.0001, respectively).

**S10 Fig. Geographical distribution of 328 wild isolates.** Map of wild isolates. Strains are colored by the variation haplotype at sqst-5 (Wild-type: grey, N2: orange, CB4856: navy, Deletion: magenta, Other putative structural variation: light pink). Strains with the deletion are labeled.

**S11 Fig. Genome-wide association (GWA) mapping identifies eight QTL across four traits in response to high zinc. A)** Normalized residual phenotype (y-axis) of 81 wild isolates (x-axis) in response to zinc supplementation. Strains are colored by the parental strains N2 (orange) and CB4856 (blue) or by the *sqst-5* variation (Deletion: magenta, other variation: light pink) **B)** GWA results are shown. Genomic position (x-axis) is plotted against the *-log10(p)* value (y-axis) for each SNV. SNVs are colored pink if they pass the genome-wide eigen-decomposition significance threshold designated by the dotted grey line. The solid grey line represents the more stringent Bonferroni significance threshold. The genomic regions of interest that pass the significance threshold are highlighted by blue rectangles. **C)** For each QTL, the normalized residual phenotype (y-axis) of strains split by genotype at the peak marker (x-axis) are plotted as Tukey box plots. Each point corresponds to a wild isolate strain. Strains with the N2 reference allele are colored grey, and strains with an alternative allele are colored navy.

**S12 Fig. Linkage mapping summary for drug-response traits in response to four heavy metals.** Genomic positions (x-axis) of all QTL identified from linkage mapping are shown for each drug-trait (y-axis). Each QTL is plotted as a triangle at the genomic location of the peak marker and a line that represents the 95% confidence interval. QTL with right side up triangles have a negative effect size (N2 allele is resistant), and QTL with upside down triangles have a positive effect size (CB4856 allele is resistant). QTL are colored by the logarithm of the odds (LOD) score, increasing in significance from purple to green to yellow.

**S1 File. Dose response phenotype data.** Processed phenotype data from zinc dose response (standard HTA)

**S2 File. Zinc response heritability.** Phenotypic values and used to calculate heritability and calculated heritabilities for all four zinc response traits (standard HTA)

**S3 File. RIAIL phenotype data.** Phenotypic values for all 121 set 1 RIAILs, 253 set 2 RIAILs, and parent strains (N2 and CB4856) in response to zinc (standard HTA)

**S4 File. Linkage mapping results.** Linkage mapping LOD scores at 13,003 genomic markers for all four zinc-response traits with the set 2 RIAILs

**S5 File. Summary of two-dimensional genome scan.** Summary of the scan2 object containing data from the two-dimensional genome scan with animal optical density (median.EXT) in zinc

**S6 File. List of zinc-related genes.** List of all previously known zinc genes (zinc transporters and hits from mutant screens), their location in the genome, and if they have variation in CB4856

**S7 File. NIL sequence data.** VCF from the whole-genome sequencing for all the NILs in this study

**S8 File. NIL genotype data.** Simplified genotypes of the NILs in the study

**S9 File. NIL phenotype data.** Raw pruned phenotypes for the NILs on chromosomes III, IV, V, and X (standard HTA).

**S10 File. Statistical significance for NIL and CRISPR assays.** Pairwise statistical significance for all strains and high-throughput assays

**S11 File. ChrV NIL breakup phenotype data.** Raw pruned phenotypes for the NILs used to break up the QTL interval on chromosome V (standard HTA)

**S12 File. Modified HTA dose response phenotype data.** Raw pruned phenotypes for the parental dose response with the modified HTA

**S13 File. ChrIII dominance assay phenotype data.** Raw pruned phenotypes for the chromosome III dominance assay (modified HTA)

**S14 File. Genes in the chrIII QTL.** List of all genes in the chromosome III interval, their functional descriptions and GO annotations, and if they have variation in CB4856

**S15 File. Expression QTL mapping results for *sqst-5*.** Linkage mapping results for the *sqst-5* expression data obtained with the set 1 RIAILs

**S16 File. Mediation estimates for chrIII QTL.** Mediation estimates for the chromosome III QTL, including *sqst-5*

**S17 File. N2 and CB4856 *sqst-5* deletion phenotype data.** Raw pruned phenotypes for the *sqst-5* deletions in the parental backgrounds (modified HTA)

**S18 File. NIL *sqst-5* deletion phenotype data.** Raw pruned phenotypes for the *sqst-5* deletions in the NIL (ECA859) background (modified HTA)

**S19 File. Reciprocal hemizygosity for *sqst-5* phenotype data.** Raw pruned phenotypes for the reciprocal hemizygosity assay using the *sqst-5* deletions in the NIL background (modified HTA)

**S20 File. Sequence of *sqst-5* in N2 and CB4856.** Raw sequence of *sqst-5* for both N2 and CB4856

**S21 File. *sqst-5* gene alignment.** Nucleotide alignment of *sqst-5* for N2 and CB4856

**S22 File. SQST-5 protein alignment.** Protein alignment of SQST-5 for N2 and CB4856

**S23 File. Structural variation in *sqst-5* in 328 wild isolates.** List of all 328 wild isolates, their isolation location, and extent of structural variation in *sqst-5*

**S24 File. *sqst-5* phylogenetic tree.** Neighbor-joining tree for all 328 wild isolates using variants near *sqst-5*

**S25 File. Wild isolate phenotype data.** Residual phenotypic values for all 81 wild isolates in response to zinc (standard HTA)

**S26 File. GWA mapping results.** GWA mapping significance values for all markers across the genome for all four zinc-response traits

**S27 File. Heavy metals linkage mapping results.** Linkage mapping LOD scores at 13,003 genomic markers for all four metal-response traits with the set 2 RIAILs for four heavy metals

